# A compact, ultrahigh-density headstage with high-fidelity hybrid integration for large-scale deep-brain opto-electrophysiology

**DOI:** 10.1101/2023.10.02.560174

**Authors:** Sungjin Oh, Kanghwan Kim, Jose Roberto Lopez Ruiz, Nathan Slager, Eunah Ko, Mihály Vöröslakos, Vittorino Lanzio, Hyunsoo Song, Sung-Yun Park, Euisik Yoon

**Author notes:** Corresponding authors: Sung-Yun Park and Euisik Yoon.

## Abstract

Recent neuroscientific research seeks to comprehend the sophisticated deep-brain networks of neural circuits consisting of large scale neuronal ensembles across multiple brain regions. An ideal way to unveil the complex connectome might be stimulating individual neurons with high spatial resolution in a broad range of brain, while seamlessly monitoring the correspondent neuronal activities. Optogenetics is known as a key technology to enable such a goal thanks to its high spatial and temporal selectivity in neuromodulation. Existing silicon probe technologies have been able to partially achieve such a goal by recording broad region of brain activities through multiple electrodes per shank, but those cannot complete perfect coverage due to the limited channel counts for the optogenetic stimulation. Here, we present an high-channel-count optogenetic system with simultaneous 256 recoding and 128 optogenetic stimulation sites, exhibiting the highest channel density ever reported, enabled by a flexible polyimide cable-based hybrid-integration of a low-stimulation-artifact micro-LED (µLED) opto-electrode with a low-power and -noise, area-efficient CMOS interfacing integrated-circuit (IC). The presented optogenetic system provides 256-neuron-size electrodes (11 × 15 µm^2^) with a 40 µm inter-electrode pitch for high spatial oversampling in recording and 128-soma-size µLEDs (8 × 11 µm^2^) with a 20 µm inter-LED pitch for single-cell resolution in stimulation, resulting in a vertical span of 640 µm and a horizontal span of 2,100 µm with a total 8 shanks. For versatility in optogenetics-based experiments from small rodents to primates with user-preferable settings, the system base that provides programmability of recording and stimulation parameters and rest of signal processing, such as filtering, digitization, and data transmission including serial peripheral interface (SPI) has also been designed within small area of 23.8 × 28.8 mm^2^ with only 3.5-gram weight, resulting in the highest channel density both in size (0.56 channels/mm^2^) and weight (109.71 channels/gram) among the state-of-the-art optogenetics-based neuromodulation systems. To verify the system operation *in vivo*, a compact optogenetics headstage has been also fabricated. Using the prepared optogenetic headstage, 169 isolated neurons have been observed with various stimulation intensities. The results offered in this article indicate that the presented hybrid integrated ultrahigh-density, high-channel-count headstage can be used to realize the massive-scale in-depth brain studies with optogenetics.

## Introduction

Monitoring complex neural activities is essential to understand the brain architecture. Optogenetics is a powerful tool to observe neural activities and explore brain structures precisely. By harnessing genetically modified neurons, the optogenetics can selectively excite or inhibit the activity of the target neurons in a single cell level, unlike other types of prevalent neural stimulation such as electrical, magnetic, or ultrasonic stimulations that blindly affect not-intended regions of brain [1–4]. Thanks to such high selectivity, the optogenetics has been widely accepted since its inception in diverse *in vivo* animal experiments and *in vitro* cellular assays, especially in those where highly selective neuromodulations are required [2–12].

Given that an ideal way to fully understand how the brain implement cognitive and behavioral functions is monitoring wholistic neuronal ensembles across multiple brain regions with a single-cell resolution, the optogenetics modulating large number of neurons with a broad spatial coverage through neuron-sized optical sources could be ideal. Advances in microfabrication technologies led to invention of high-density micro-LED (µLED) optoelectrodes, expanding the application of the optogenetics to extensive investigation of the brain circuits with a single-cell resolution. For instance, various neuronal circuit dynamics in hippocampus, such as excitatory/inhibitory networks, paradoxical short- and long-range interactions, and functional connections among three different types of neurons, were observed through optical perturbations using the high-density µLED optoelectrodes [13–18]. Furthermore, the µLED optoelctrodes may also enable studying striatum, amygdala, and other architecturally complex regions in deep-brain where large-scale high-density illumination is necessary to untangle microcircuits and their behavioral outcomes [19–21]. Despite this improvement through the µLED optoelectrodes, lack of a compact and complete large-scale opto-electrophysiology interface still prevents the optimal use of the optogenetics in diverse animal experiments for understanding complex neuronal circuits across broad brain regions. Therefore, it is crucial to implement a more advanced optogenetic system that can provide broader spatial coverage through higher channel counts with a smaller form-factor.

Recent advances in microfabrication techniques have enabled highly dense arrays of recording sites via silicon-based microelectrodes. To enhance the recording quality and to relax interconnection issue in the dense array, monolithic integration of complementary metal oxide semiconductor (CMOS) integrated circuit (IC) and microelectrode has been explored by adopting the advanced CMOS technologies [22–27]. However, to realize large scale optogenetics, multiple synchronized optical stimulation also has to be implemented in the monolithically integrated platform. The current monolithic approach fails to do so, resulting in large, dense recording but spatially sparse stimulation due to the difficulties in integration of miniaturized optical sources, such as μLEDs or waveguides in the existing microelectrodes. Another approach toward large scale optogenetics is to develop a full-custom fabrication technology without resort to CMOS processes incompatible with optical sources. Several biosensors have been developed using organic thin film transistors fabricated in full-custom methods [28–32], those can provide higher level of integration of optical sources than silicon-based microelectrodes. However, poorer electrical performance of the custom-fabricated electronics results in lower bandwidth and higher noise per power, and larger area consumption than silicon-based integration, leaving them way behind the silicon-based microelectrodes in scaling up toward large optogenetic systems. The third approach is hybrid integration where each component for system is separately fabricated, and then combined through customized packaging techniques. Since each component solely depends on each dedicated fabrication process, this approach will be able to allow implementation of high-density optogenetic systems without the conflicts appeared in the monolithic integrations. Up to now, various hybrid integration techniques that can be potentially adoptable for large-scale optogenetic systems have been presented in prior arts [33–43]: using bulky interconnections through mechanical bonding or commercial-off-the-shelf (COTS) components [33–41], or flip-chip bonding of the CMOS IC to microelectrodes [42, 43]. However, none of them has implemented a large-scale optogenetic headstage with a compact form factor that fits into small size living animals with simple, reliable, and repeatable integration. Understandably, there is no existing large-scale optogenetic systems up to date.

In this article, we present an ultrahigh-density, *i.e.*, a large-scale within small form factor optogenetic system through the high-fidelity hybrid integration. The presented optogenetic system has broad brain coverage over 0.6-mm vertical and 2-mm lateral spans with simultaneous 256 neural recording and 128 optical stimulation sites through a multi-shank μLED optoelectrode and CMOS interface IC, which is the largest scale optogenetic system ever reported. For our integration, a polyimide-based flexible interposing cable has been fabricated and used as a hybrid integration vehicle for chip-to-μLED optoelectrode interconnection and a two-step ball-bumping process that guarantees high yield have been employed. Thus, our optogenetic system can achieves the highest channel density in size of 0.56 channels/mm^2^ and weight of 109.71 channels/gram even when considering the entire headstage for *in vivo* that includes some redundancy for easy handling in experiments. Besides, the interposing cable enables only a small part of our optogenetic system to be implanted in a brain rather than the entire device, relieving surgical difficulties, increasing mechanical robustness, and reducing risk of tissue damage due to the heat flux emitted by the active electronics in the CMOS IC.

### Overview of the hybrid integrated ultrahigh-density optogenetics system

A conceptual diagram of the proposed optogenetic system-based headstage implanted in a mouse brain is shown in Fig. 1(a). The headstage hybrid-integrates an 8-shank high-density low-stimulation-artifact μLED optoelectrode and a high-channel-count CMOS interface IC through a polymer-based flexible, long-interposing cable, covering a broad range of brain both in depth and width. This headstage configuration also can relax many practical issues in experiments. First, it can minimize the volume of the implanted part to only the optoelectrode and a part of the cable with sub-micron thickness, while leaving other system components such as CMOS IC and interconnect out of the implantation; thus, significantly relieves the surgical damage compared to the full-system implant. In addition, it can keep the brain temperature stable during the experiment because the CMOS IC and other electronic components, which has a potential to heat up the surroundings through its power dissipation, are located far outside from the brain. It was well-verified that the μLEDs fabricated on the optoelectrode emit the negligible amount of heat [18]. Moreover, the flexible cable softens the implanted device, and eventually enables robust neuromodulation in a long-term period [44–46]. Finally, the volume and weight of the headstage can be significantly reduced since the sub-micron-scale interposer replaces the bulky off-the-shelf connectors that were widely used in the previous works [34–40].

**Fig. 1.**
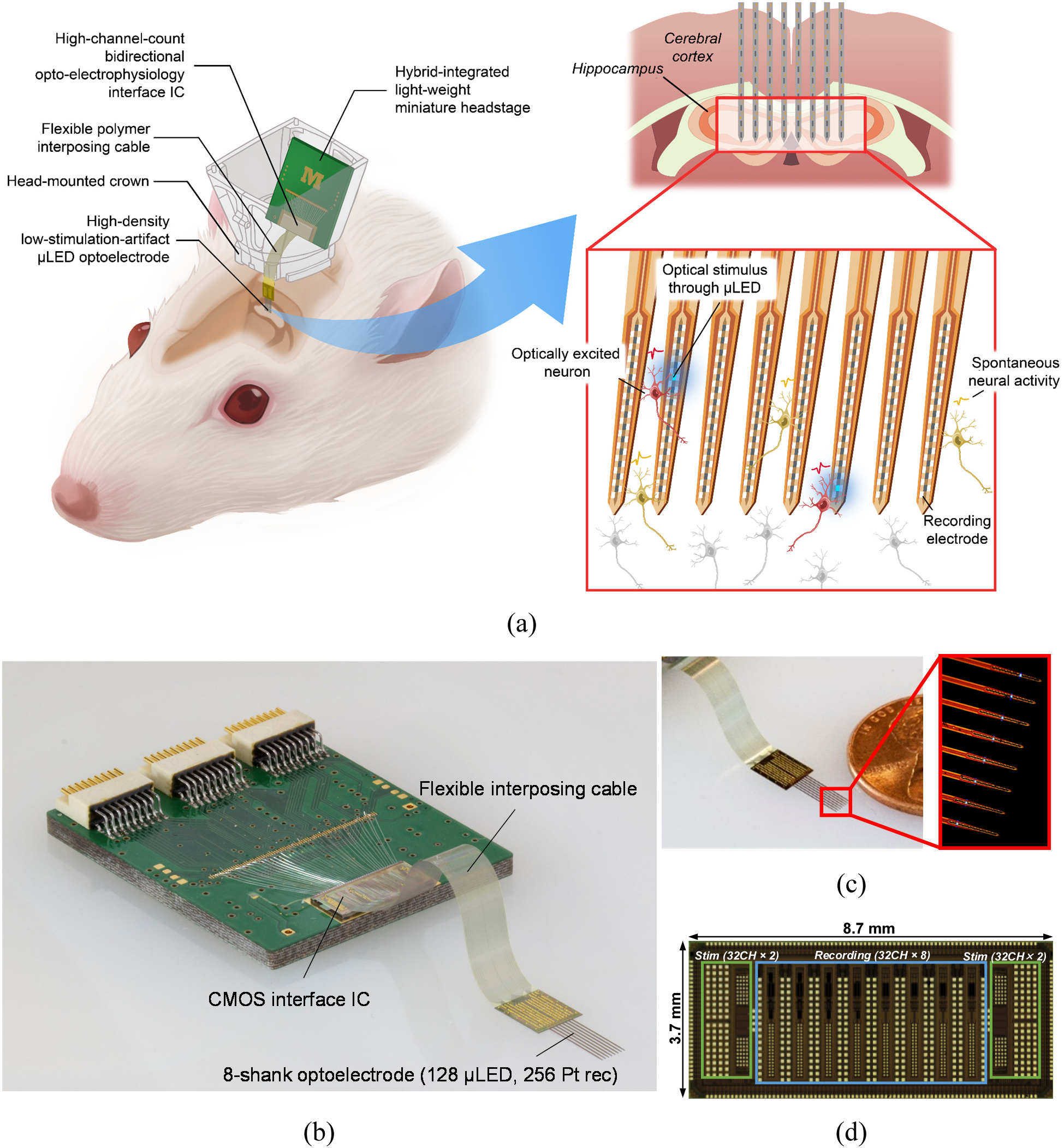
(a) Concept diagram of the hybrid integrated headstage for large-scale deep-brain opto-electrophysiology. (b) Photo of the fully integrated headstage composed of the high-density low-stimulation-artifact µLED optoelectrode, high-channel-count CMOS interface IC, and flexible polyimide interposing cable. (c) Microphotograph of the 8-shank high-density low-stimulation-artifact optoelectrode. (d) Microphotograph of the high-channel-count CMOS interface IC.

Fig. 1(b) shows the photograph of the proposed optogenetics system-based headstage. Note that the volume is mostly constrained by the COTS connectors and the headstage has been intentionally designed large for easy handling with finger tips. All the headstage electronics including the CMOS interface IC were integrated on a printed circuit board (PCB) of 23.8 × 28.8 mm^2^, which is still small enough to be mounted on a mouse head. The weight of the fully assembled headstage is only 3.5-gram, sufficiently light to be used for the *in vivo* experiments with a small rodent, even considering the strict restriction of the device weight, ≤10% of the body weight, to avoid physical stress to the animal from heavy over-hanging fixture in its head [47]. Fig. 1(c) and (d) show the microphotographs of the microfabricated components, 8-shank high-density low-stimulation-artifact μLED optoelectrode and high-channel-count CMOS interface IC, respectively.

### Hybrid integration of headstage components

To implement the headstage with the ultrahigh density, it is vital to choose the optimal procedure of integrating the microfabricated components with high yield. Since the total 128 stimulation channels as well as 256 recording channels should be configured within miniaturized areas on the optoelectrode, the interposing cable, and the CMOS interface IC, the integration procedure must reliably interconnect those components via small pad-to-pad pitches less than 100 μm. In order to provide the near perfect bonding yield with a fine pad-to-pad pitch less than 100 μm, we chose ball-bump bonding that is well-known for the ignorable defect rate (< 0.0001 %) in the packaging industry [48].

The detailed integration procedures of the proposed headstage are shown in Fig. 2(a). We employ a two-step ball-bump bonding process: i) bonding of the interposer to the CMOS interface IC and ii) bonding of the interposer to the optoeletrode. The data/power connectors (PZN-18-AA, *Omnetics*, MN, USA), used to link the interface IC and a back-end control system, first are soldered on the headstage PCB. Then, the CMOS interface IC is glued on the large-area gold-plated ground pad on the PCB by using a conductive adhesive, in order to minimize the circuit ground noise. After the flexible interposing cable is aligned and bonded to the CMOS interface IC through the 1^st^-ball-bump bonding, the signal/power pads of the interface IC are wire-bonded to the headstage PCB. Then, the optoelectrode is integrated with the flexible interposing cable through the 2^nd^-ball-bump bonding. Finally, the peripheral electronics, such as voltage regulators and reference generator (those are optional), and other necessary passives are soldered on the backside of the headstage PCB. The schematic diagram of the fully assembled headstage is shown in Fig. 2(b). Since the circuit power and reference signals can also be supplied through the connectors according to the user preference, the voltage regulators and reference generator, much larger and heavier than the other passive components, can be removed to further reduce the headstage weight to less than 3.5-gram.

**Fig. 2.**
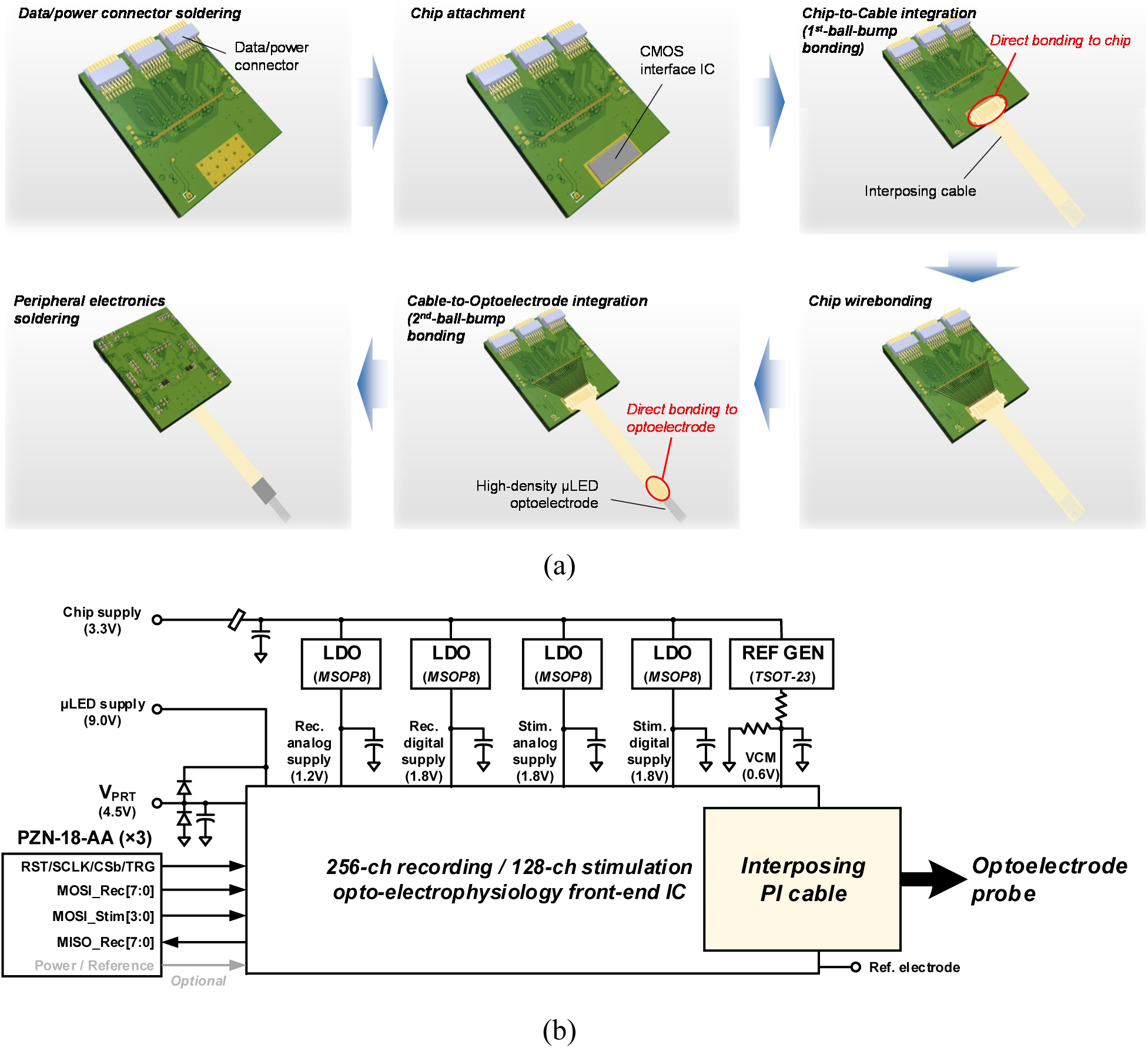
(a) Hybrid integration process of the headstage: 1) Soldering data/power connectors on the bare headstage PCB. 2) Attaching the CMOS interface IC on the headstage PCB using the conductive adhesive. 3) Integration of the flexible interposing cable to the interface IC (1^st^-ball-bump bonding). 4) Wire-bonding of the interface IC to the headstage PCB. 5) Integration of the high-density μLED optoelectrode to the flexible interposing cable (2^nd^-ball-bump bonding). 6) Soldering peripheral electronics backside. (b) Schematic of the fully integrated headstage.

### High-density low-stimulation-artifact μLED optoelectrode and flexible interposing cable

At the interface between the headstage and the animal’s brain is the μLED optoelectrode [18, 49, 50]. The μLED optoelectrode is capable of recording neuronal activities from neurons in its surroundings at single-cell resolution while providing optical stimulation at a resolution comparable to the recording resolution. The gallium-nitride/indium-gallium-nitride (GaN/InGaN) μLEDs on the optoelectrode emit blue light, whose peak wavelength is approximately 465 nm, from their top surfaces. Photons generated from each μLED deliver optical energy to a roughly hemispherical volume surrounding the μLED, and this volume overlaps with the volume from which the adjacent platinum (Pt) electrodes pick up electrical signals [18], as depicted in Fig. 1. Thanks to the high-density, monolithic integration of stimulating μLEDs and signal recording sites, the μLED optoelectrode enables the precise delivery of optical stimulation to the target brain volume.

High-density, low-stimulation-artifact μLED optoelectrodes were fabricated utilizing miniSTAR [49] and hectoSTAR [18] technologies. More specifically, GaN/InGaN LED stack was carefully deposited on a heavily-boron-doped silicon (Si) substrate [49] ultra-thin-and-narrow (0.1-μm thick and 0.7-μm wide) metal lines were carefully stacked up in a multi-metal-layer configuration [18], and a grounded shielding layer was included as a part of the multi-metal-layer stack [49]. Snapshots of the cross-section of a high-density, low-stimulation-artifact μLED optoelectrode after a few key steps of the fabrication process are shown in Fig. 3(a).

**Fig. 3.**
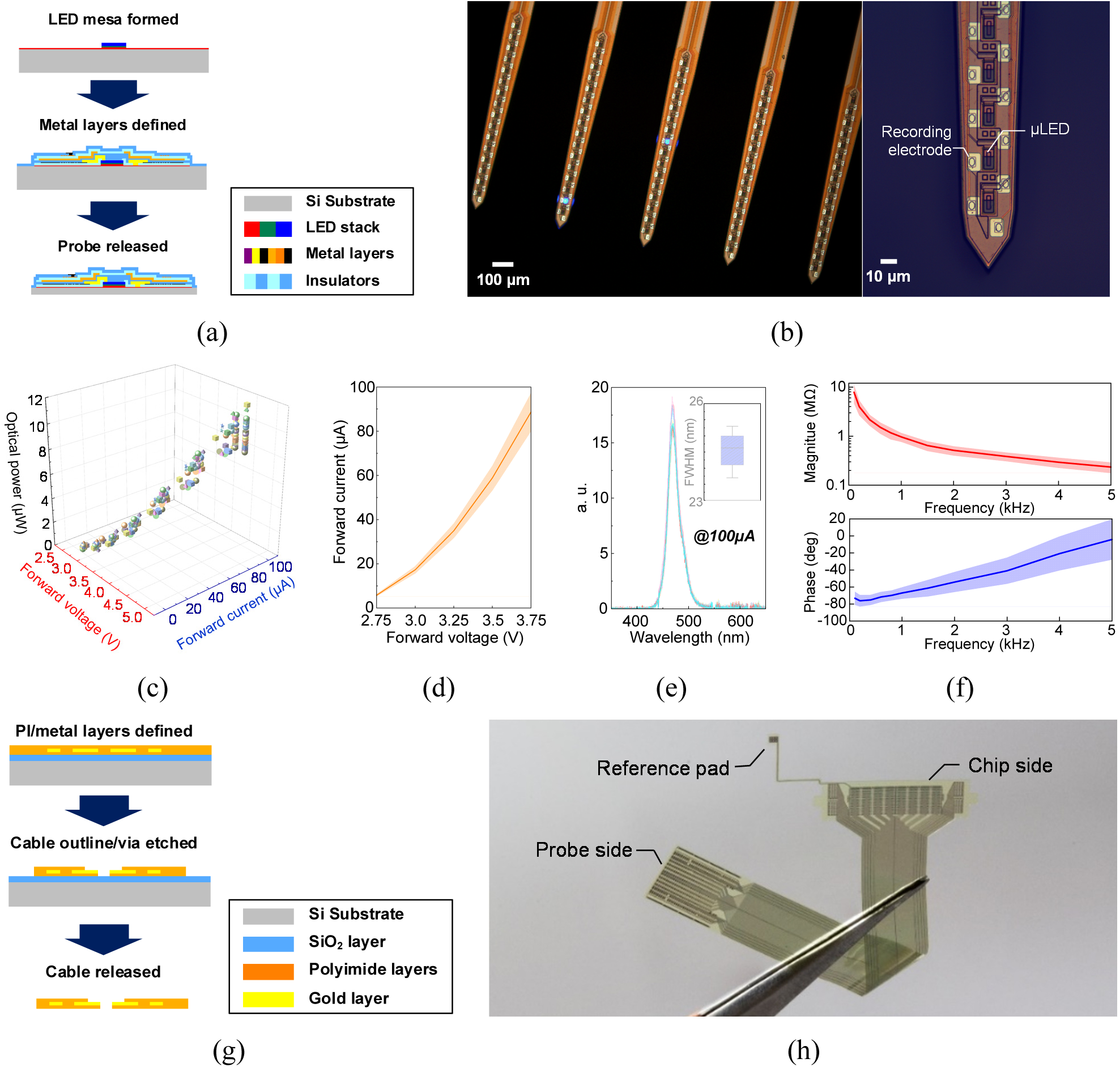
(a) Fabrication process of the high-density low-stimulation-artifact μLED optoelectrode. (b) Microphotograph of the fabricated 8-shank high-density low-stimulation-artifact optoelectrode. (c) Measured optical power versus forward voltage and current of the fabricated μLEDs (n = 40). (d) Measured I-V characteristics of the fabricated μLEDs (n = 40). (e) Measured optical power spectrum of the fabricated μLEDs (n = 10). (f) Measured impedance magnitude and phase of the fabricated Pt recording electrodes (n = 40). (g) Fabrication process of the flexible polyimide interposing cable. (h) Photo of the fabricated flexible polyimide interposing cable.

Fabricated μLED optoelectrode features 16 GaN/InGaN μLEDs and 32 platinum (Pt) electrodes on each of its shank, each of which is integrated at 20-μm and 40-μm vertical pitch (or at 50 mm^-1^ and 25 mm^-1^ linear density), respectively. The total 8 shanks are configured in the single optoelectrode; thus, it includes 128 μLEDs and 256 Pt electrodes within a deep vertical span of 640 µm and a wide horizontal span of 2,100 µm. Fig. 3(b) shows a close-up microphotograph of the shanks on the fabricated μLED optoelectrode. Each 8 × 11 μm^2^ μLED on the optoelectrode generates light with optical power up to 9.55 μW (σ = 0.93 μW, *n* = 40) with as low as 100 μA of current, and the 11 × 15 μm^2^ Pt electrodes have the impedance of 0.96 MΩ (−67.03 °) at 1 kHz (σ = 0.19 MΩ and 6.61°, *n* = 40). Fig. 3(c)−(f) shows the measured optical and electrical characteristics of 40 representative μLEDs and Pt electrodes. The characteristics of the μLEDs and electrodes suggest that they can provide a desirable near-single-cellular stimulation-and-record capability [18].

The flexible interposing cable is another important component of the headstage that sits between the optoelectrode and the interface CMOS IC, providing mechanical and electrical connections between the two components. In order to reliably provide long-lasting connections for total 384 signals between the IC and the probe, the interposer is required not only to be sufficiently compliant for the ease of handling and mechanical stability but also to be robust for the resistance to the wet and ion-rich surgical environment. Therefore, a flexible interconnect technique whose reliability has been proven over more than a decade in multiple applications including biomedical applications, polyimide interconnect cable (also referred to as microflex interconnect [51]) was utilized. The interposing cable was fabricated using a two-mask process presented in [52]. In the process, a metal layer serves as both the conductor and the hard mask for the etching of a via hole in the polyimide layer. Fig. 3(g) shows snapshots of an imaginary cross-section of a cable after a few key fabrication steps. As shown in Fig. 3(h), the cable has arrays of ball-bumping pads (with vias at the center) on both of its ends, whose positions are designed to be aligned to the center of the exposed gold and aluminum bonding pads on the optoelectrode and the interface CMOS IC. An additional short branch is attached on the chip side of the cable so that the reference potential, which is wired through the PCB to the animal ground, can be wired through the branch. This branch is connected to the PCB via direct ball-bump bonding on an electroless nickel immersion gold (ENIG) pad on the printed circuit board. Fabricated using relatively thick (400 nm) and wide (2 µm) gold lines, the cable adds < 1 kΩ resistance to each signal route, which is far less than the electrode impedance, and therefore minimally affects the electrical performance of the proposed optogenetics system.

### High-channel-count CMOS IC for optogenetics system

To implement bidirectional control of optogenetics using the fabricated high-density low-stimulation-artifact μLED optoelectrode, the CMOS interface IC should accommodate the large number of stimulation and recording channels. Compact and energy-efficient architectures are necessary to implement the interface IC with such the large number of channels. In addition to such design requirements, the interface IC has to grant high-precision in both the optical stimulation control and electrical recording resolution to the optogenetics system to enable quality-guaranteed neuromodulations. Fig. 4(a) shows the system architecture of the proposed high-channel-count interface CMOS IC. The interface IC can support 128 optical stimulation channels with 8-bit illumination precision (*i.e.*, separate 256 steps), as well as 256 broadband neural recording channels with 10-bit analog-to-digital conversion precision. Both the stimulation and recording circuits were implemented through a modular architecture: each module controls 32 channels either in stimulation or recording. The number of channels can be easily scaled-up without complicated design modifications thanks to the modular architecture. A dedicated serial peripheral interface (SPI) is used to digitally control the interface IC according to commands from the back-end control system.

**Fig. 4.**
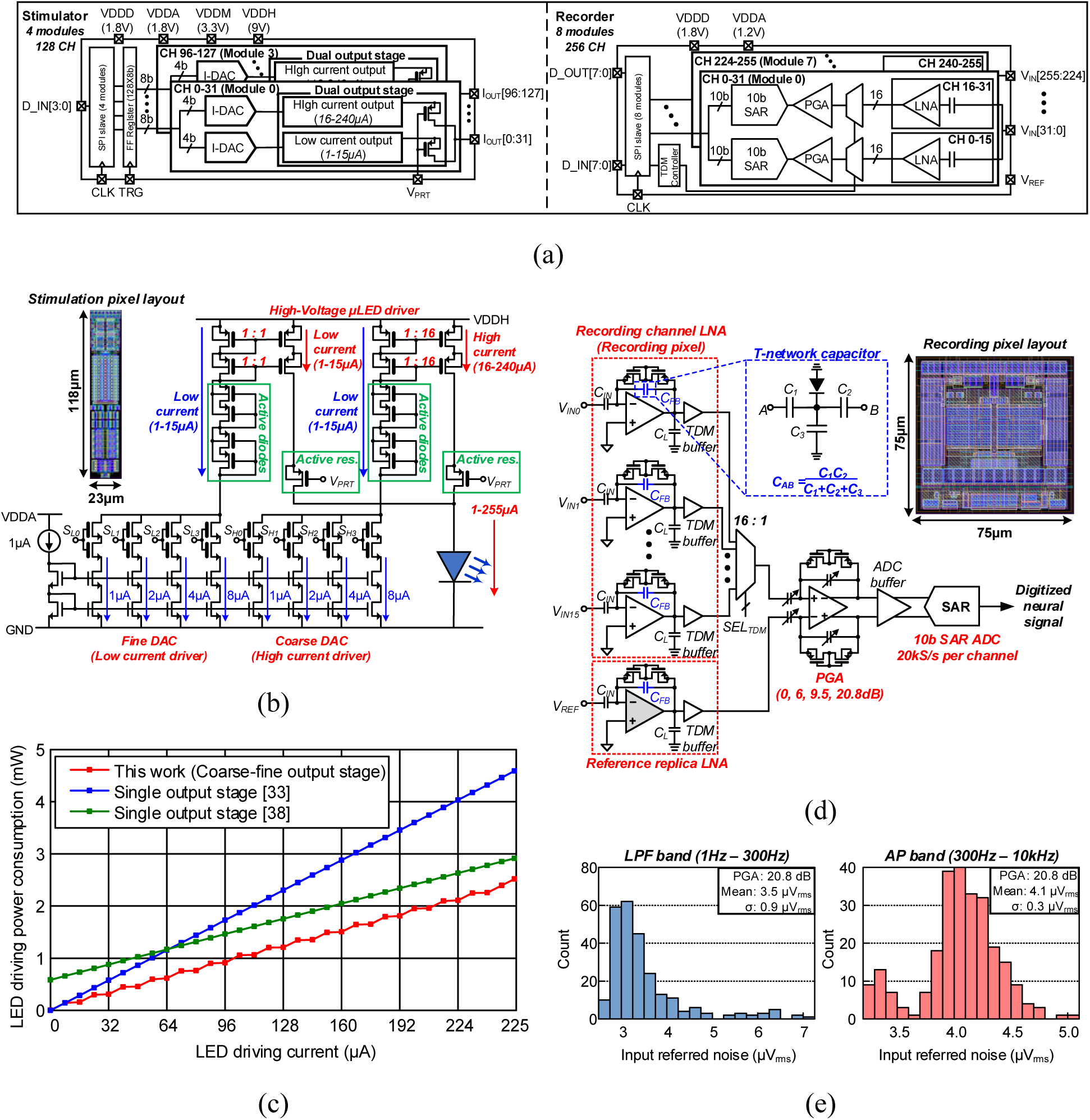
(a) System architecture of the high-channel-count CMOS interface IC. (b) Schematic and layout of the optical stimulation circuit. (c) Saving of the LED driving power through the coarse-fine output stage design. (d) Schematic and layout of the broadband neural recording circuit. (e) Measured input referred noises of the fabricated recording circuit.

Fig. 4(b) shows the schematic of the stimulation control circuit. To linearly adjust the optical stimulus intensity and duration, we chose the current-controlled stimulation that inherits a higher linearity than other approaches, such as voltage-controlled or switched-capacitor stimulation [39, 53]. The output stages are segmented with coarse and fine current generation blocks. Two 4-bit binary current-steering digital-to-analog converters (DACs) with 1 µA LSB are used to adjust the stimulation current amplitude. The fine output stage generates the low amplitude stimulation current in 1 – 15 µA by copying the fine DAC output current (1 – 15 µA) without amplification, as similar to the conventional design using a single output stage [33]. On the other hand, the coarse output stage copies the coarse DAC output current (1 – 15 µA) with ×16 amplification so that it generates the large amplitude stimulation current up to 240 μA. Therefore, the total current amplitude that should be copied from the current-steering DAC to generate the maximum stimulation current of 255 µA is only 30 µA, reduced by 88.2 % compared to the conventional design using the single output stage without amplification [33]. As shown in Fig. 4(c), this results in >34.1 % save in the average power consumption than the previous works using the single output stage [33, 38]. The size of the current-steering DAC can also be reduced by adopting the coarse-fine output stages. Since the size of transistors implementing a current-steering DAC is proportional to its output current amplitude, the current steering DAC dedicated to the coarse-output stage can be implemented in a 93.8 % smaller area than the counterpart used for the single output stage [33]. To cope with the various forward voltages of the fabricated μLEDs, the output stages should drive the stimulation current with a sufficiently high voltage. Therefore, the stimulation circuit uses a 9-V supply. Since the CMOS process used in this work cannot accommodate the supply voltage higher than 3.3 V, active diodes and resistors are added to accommodate the 9-V supply. The high-temperature operating life test at 70 °C verified that those additional components increased the life time of the stimulation circuit operating with the given stimulation parameters (255 μA at 4 Hz frequency and 50% duty-cycle) at the room temperature from ∼30 days to > 4 years [54]. Each stimulation channel circuit is integrated in a small area of 23 × 118 μm^2^ and consumes quiescent power of 2.2 μW.

The recording circuit is implemented with the architecture presented in our prior work [42]. Fig. 4(d) shows the schematic of the half-module of the recording circuit. The neural signals are amplified and digitized through a signal chain: a low-noise amplifier (LNA), a programmable gain amplifier (PGA), and an analog-to-digital converter (ADC). The LNA amplifies the tiny neural signals with a large voltage gain of 42 dB, as well as filters out-of-band noise (>10 kHz) and electrode DC offset drifts (<1 Hz) through capacitive-coupled inputs [55]. The PGA is added to prevent possible circuit saturation caused by deviations in the recording environment, *e.g.,* recording location and electrode impedance [56]. Finally, the ADC digitizes the magnified neural signals for robust data transmission to the back-end control system. To implement the signal chain within the limited power and area, the PGA and ADC are shared by the 16 LNAs through time-division multiplexing [42, 57]. For the smaller size of each recording channel circuit, it is essential to minimize the capacitors used in the LNA. Two design techniques are adopted to integrate the smaller capacitors. First, the LNA is implemented as a single-ended amplifier with a single capacitive-coupled input, which reduces the capacitance by half compared to the conventional design with a pair of the capacitive-coupled inputs [42]. Second, a T-network capacitor architecture is adopted to implement the feedback capacitance (*C_FB_*) lower than the process-minimum capacitance value [58]; thus, eventually reducing the input capacitance (*C_IN_*) by 29.8 % compared to the design without the T-network. The PGA provides the variable amplification gain from 0 to 20 dB. The successive approximation register (SAR) ADC digitizes the amplified neural signals with 10-bit resolution. Since the SAR ADC is shared by the 16 recording channels, the conversion rate is set at 320 kS/s to implement the per-channel Nyquist rate of 20 kS/s. The whole signal chain achieves the sufficiently low input-referred noise (IRN) to acquire subcortical neural signals: 3.5 – 5.1 μV_rms_ in the local field potential (LFP) band (1 – 300 Hz), and 4.1 – 8.1 μV_rms_ in the action potential (AP) band (300 Hz – 10 kHz). Each recording channel circuit is integrated within a miniaturized area of 75 × 75 μm^2^ and consumes 29.6 μW.

### Back-end control system

Both hardware and software systems were implemented in the back-end module to conveniently control the headstage. The back-end system uses the concise signal chain as shown in Fig. S4(a). The SPI configures data transactions between the CMOS interface IC on the headstage and the microcontroller implemented with a complex programmable logic device (CPGA). The CPLD serializes the stimulation/recording circuit control commands generated by the customized software programs, while deserializing the obtained neural signals. For software-to-hardware data conversion or vise-versa, a commercial digital I/O device (PCIe-6536, *National Instruments*, TX, USA) is used. Thanks to the modular configuration of the I/O device, it is available to use multiple headstages simultaneously. As shown in Fig. S4(b), the customized software program enables modifying the optical stimulation parameters of 128 μLEDs while simultaneously monitoring the neural signals at 256 recording sites in real-time.

### *In vivo* experiments

To validate the functionality of the fabricated headstage, we performed *in vivo* experiments that monitored brain activities of a head-fixed anesthetized Thy1-ChR2-YFP mouse. The optoelectrode connected with the flexible interposing cable was implanted in the hippocampus of the mouse to measure subcortical neuronal activities, while the other part of the headstage was located outside of the mouse brain. The broad horizontal span (2,100 µm) of the fabricated 8-shank optoelectrode allows to record the dorsal hippocampus bilaterally as illustrated in Fig. 5(a). Wideband brain signals were recorded, including both slow-varying LFP band signals (< 300Hz) and high-frequency single-neuron APs (3 – 6 kHz).

**Fig. 5.**
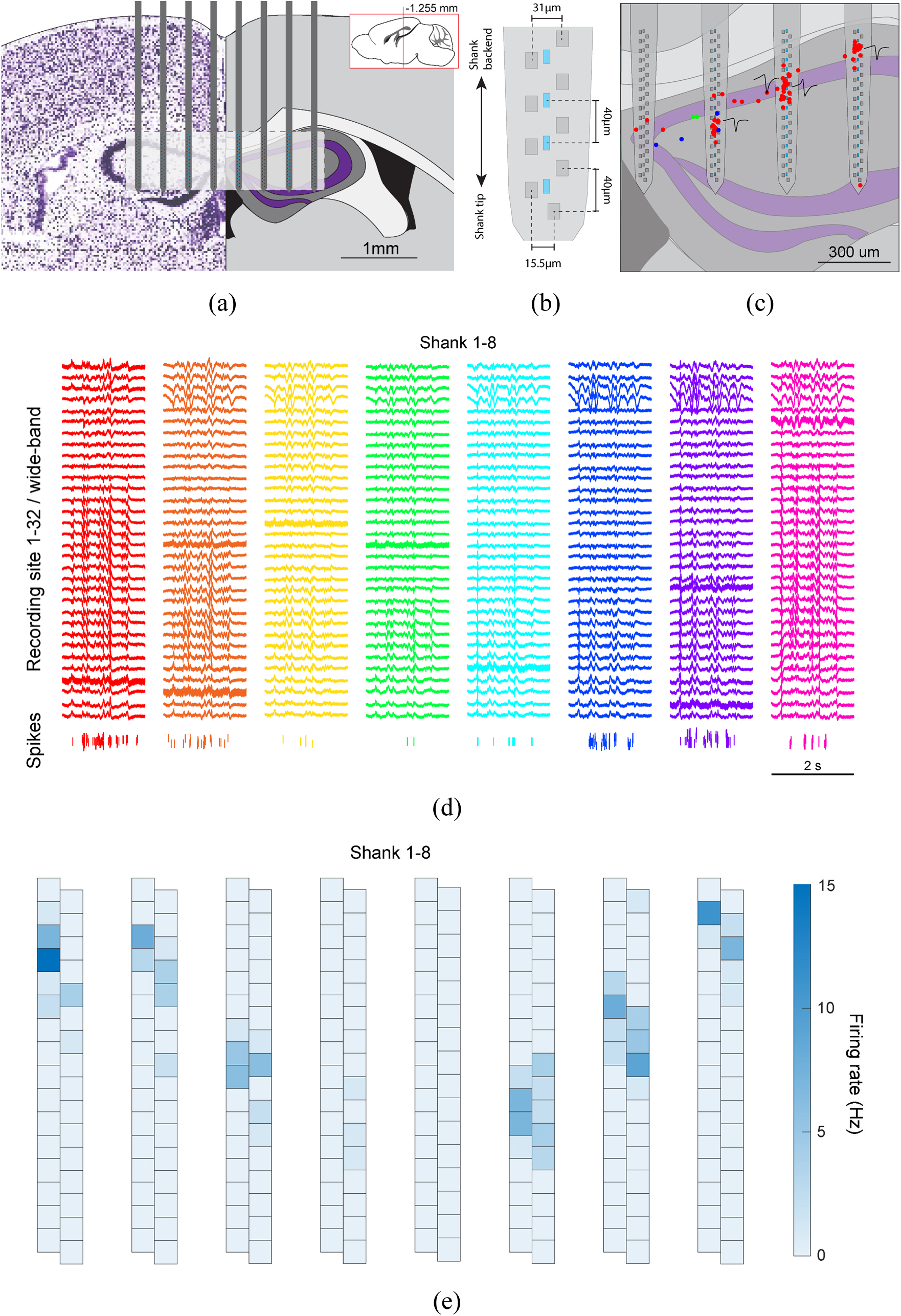
Recorded spontaneous neural activities in the anesthetized mouse dorsal hippocampus. (a) Anatomy of the bilateral hippocampus recording: Optoelectrode shanks are shown on top of the dorsal hippocampus anatomy. (b) Shank topography. (c) Estimated positions of the captured single neurons in the right hippocampus. (d) Spontaneous transient responses recorded from the dorsal hippocampus and the corresponding raster plots of the single unit activities. (e) Firing rate of the obtained single neurons.

In the first phase of the experiment, the spontaneous electrophysiology from the dorsal hippocampus was monitored with the wide spatial coverage of the fabricated headstage. With the given topography of the recording sites shown in Fig. 5(b), we were able to accurately estimate the position of the nearby neurons whose activity was captured by the headstage as shown in Fig. 5(c). The landmark LFPs from both hippocampi were also simultaneously monitored as shown in Fig. 5(d). The total of 169 well isolated neurons were recorded: 153 putative pyramidal neurons, 12 narrow interneurons, and 4 wide interneurons. The averaged spontaneous firing rate distribution across the recording sites on the implanted optoelectrode clearly reflects the anatomy of the dorsal hippocampus as shown in Fig. 5(e).

Second, in order to study the local and long-range effects of focal optical stimulation in the dorsal hippocampus, we monitored the activity of the dorsal hippocampal neurons while optically stimulating different subregions of the dorsal hippocampus architecture. After an initial 15-minute baseline period, optical stimulation was applied by repeatedly activating each µLED 100 times with 5, 10, and 20 µA. We utilized a square pulse (ON-OFF) train with a stimulation frequency of 2 Hz and a duty-cycle of 20 % (100 ms pulse width). Fig. 6(a) shows the transient neural responses including LFPs and single unit activities when the µLED in the 2^nd^-shank (S2L15: second from the top) was activated at 20 µA. The transient plot shows fast frequency oscillations in the wideband recordings at the top of the 2^nd^-shank in response to the optical stimulation. The evoked single unit activity is also shown in the bottom raster plot in Fig. 6(a). The activation of the nearest µLED to the pyramidal cell layer triggered APs not only in the nearby neurons, but also in the neurons on the adjacent shanks as shown in Fig. 6(b). The strength of the evoked responses in the adjacent shanks increased with the stimulus intensity and decayed over distance to the stimulation site as shown in Fig. S6(b). It is noteworthy that these responses from the adjacent shanks were caused by the downstream and/or upstream activation of the hippocampal circuitry, as previously reported in [17, 18]. Fig. 6(c) shows the single neuron activities at the variable stimulation intensities. As expected, the number of evoked APs gradually increased as the stimulation current was changed from 5 μA to 20 μA at the nearest µLED (S1L15). Fig. 6(d) shows the single neuron responses with the activation of the different µLEDs on the same shank. Illumination of the nearest µLED (S1L13) resulted in the lowest evoked latency (< 8 ms), suggesting a direct activation of the monitored neuron. On the other hand, the activation of the µLEDs farther away from the studied neuron resulted in the longer AP latencies (10 – 25 ms), suggesting the indirect activation through the local neuronal network. Fig. 6(e) shows the single neuron activities with various locations of the optical stimulation in the three different shanks. Illumination of the µLEDs on the 2^nd^- and 4^th^-shank failed to evoke the response from the neuron on the 3^rd^-shank, which supports that the interference between the neighboring shanks is ignorable.

**Fig. 6.**
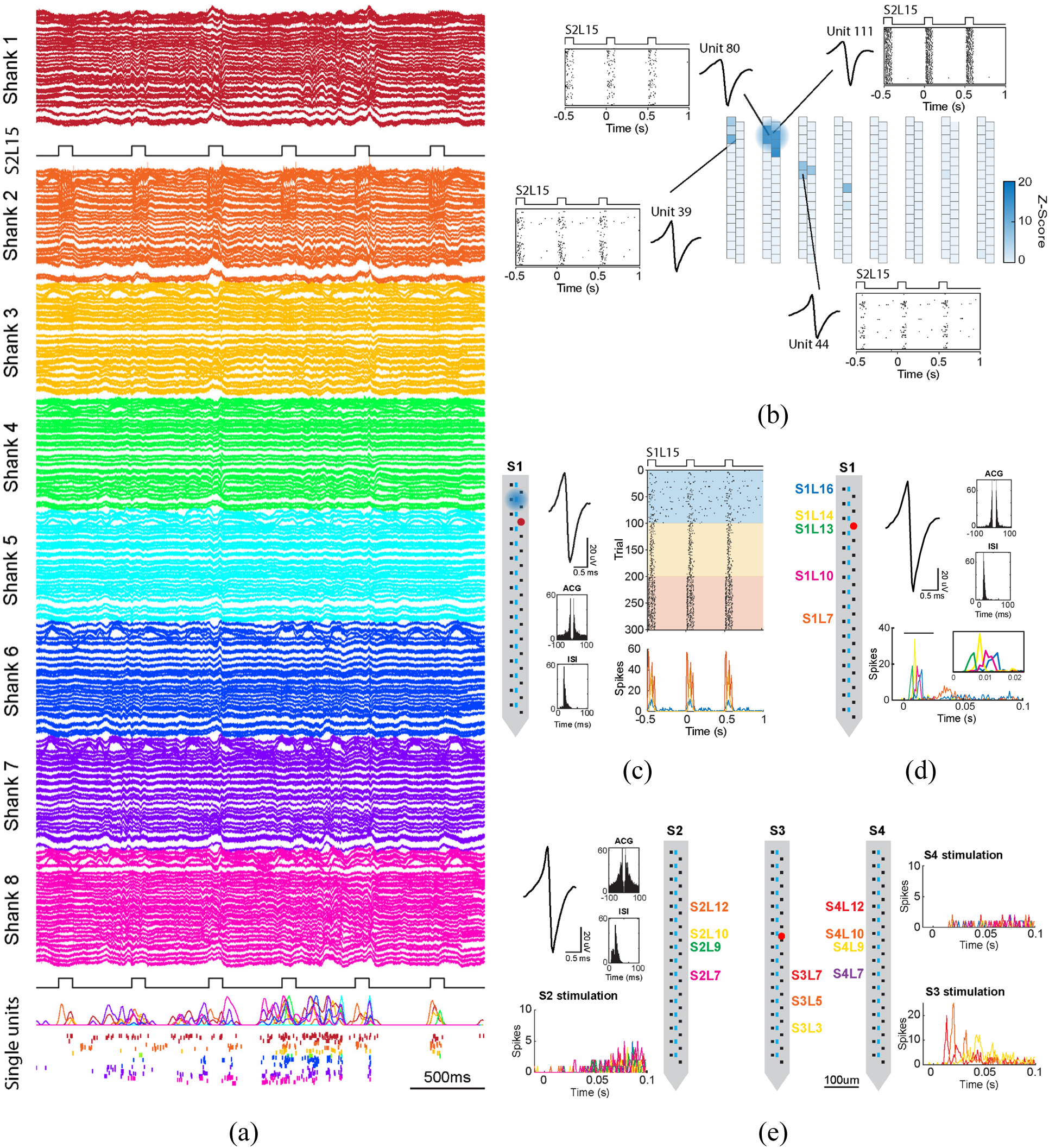
Optogenetically evoked neural activities in the anesthetized mouse dorsal hippocampus. (a) Transient responses with optical stimulation through the second-top µLED in the 2^nd^-shank (S2L15) and the corresponding raster plots and PSTHs of the single unit activities (colors are paired to each shank). (b) Spike rates in response to the optical stimulation and the raster plots of four example neurons on the stimulated and neighboring shanks. (c) Single neuron responses with the stimulation current of 5 µA (sky blue), 10 µA (yellow), and 20 µA (orange). The locations of the neuron and activated µLED are shown as the red and blue dots, respectively. (d) Single neuron responses to the stimulation from different µLEDs on the same shank (colors are paired to each µLED). The neuron was located at the red dot. (e) Single neuron responses to the optical stimulation from different µLEDs on the same or neighboring shank (colors are paired to each µLED). The neuron was located at the red dot.

### Comparison to the state-of-the-arts optogenetic systems

The presented ultrahigh-density optogenetic system is compared with the other state-of-the art optogenetic systems in Table 1. This work achieves a small form-factor opto-electrophysiology interface with the highest area- and weight-efficiencies. The compact chip-to-optoelectrode integration was achieved through the flexible polyimide interposing cable, unlike the other works used bulky and heavy off-the-shelf connectors [34, 38, 40, 59, 63]. Also, the presented work employs the custom-designed CMOS interface IC to implement the high-channel-count optical stimulation and broadband neural recording with a miniaturized size and weight, unlike majority of the prior works that used macro-packaged multiple circuit components to implement the stimulation or recording functionality [33, 38, 40, 59, 60, 61]. Although a few prior devices used a custom-designed IC, the numbers of the recording and stimulation channels that can be supported by those circuitries are ×8 to ×16 smaller than this work [34, 62, 63]. In order to quantitatively compare the area- and weight-efficiencies, we estimate the channel density by calculating the effective number of channels integrated in a unit size or weight of the headstage. This work achieves the highest channel densities in terms of the headstage size (0.56 channels/mm^2^) and weight (109.71 channels/gram) among the state-of-the-arts. Additional comparisons of the high-density low-stimulation-artifact μLED optoelectrode and high-channel-count interface IC with the other optogenetic systems are offered in Table S1 and S2.

**Table 1.** Comparison to the state-of-the-art optogenetic systems.

|  |  | **This work** | [33] | [34] | [38] | [40] | [59] | [60] | [61] | [62] | [63] |
| --- | --- | --- | --- | --- | --- | --- | --- | --- | --- | --- | --- |
| Bidirectional interface |  | **Yes** | Yes | Yes | Yes | Yes | No | No | No | No | Yes |
| Probe |  | **μLED opto electrode** | μLED opto electrode | μLED opto electrode | μLED opto electrode | Optrode w/ opticfiber | μLED needle | μLED needle | μLED needle | Planar LED array | Planar array, Tungsten needle |
| Interface circuit |  | **ASIC** | ASIC (Stim) COTS (Rec) | ASIC | ASIC (Stim) COTS (Rec) | COTS | COTS | COTS | COTS | ASIC | ASIC |
| Integration | Probe | **Ball-bump bonding** | Ball bonding | Connector | Connector | Connector | Connector | Silver epoxy | SMT | SMT | Connector |
|  | Circuit | **Wire bonding** | Wire bonding, SMT | Wire bonding | Wire bonding, SMT | SMT | SMT | SMT | SMT | Wire bonding | Wire bonding |
| No. of Ch. | Optical stimulation | **128** | 12 | 12 | 12 | 1 | 2 | 1 (4LEDs) | 1, 2, 4 | 16 | 16 |
|  | Other stimulation | - | - | - | - | - | 4 | 1 | - | - | 1 |
|  | Recording | **256** | 32 | 32 | 32 | 8 | - | - | - | - | 16 |
| Flexible implant |  | **Yes** | No | Yes | Yes | No | Yes | Yes | Yes | Yes | Yes |
| Headstage size (mm<sup>2</sup>) |  | **685.4** | 514.1 | 646 | 282.8 | 306 | 200* | 125 | 78.5 | 112 | 3500 |
| Channel size density (No. of Ch./mm<sup>2</sup>) |  | **0.56** | 0.09 | 0.07 | 0.16 | 0.03 | 0.03 | 0.04 | 0.05** | 0.14 | 0.01 |
| Headstage weight (g) |  | **3.5** | 1.9 | 4.9 | 1.37 | 4.9 | 2 | 0.22 | 0.075 | 0.5 | 17 |
| Channel weight density (No. of Ch./g) |  | **109.71** | 23.16 | 8.98 | 32.12 | 1.84 | 3 | 22.73 | 53.33*<br>* | 32 | 1.94 |

\* Estimated size from the device photo. \*\* Density at the maximum channel count.

## Conclusions

In this paper, we presented the ultrahigh-density, high-channel-count headstage that enables the massive-scale deep-brain optogenetics across the broad range of the brain. Through compactly integrating the high-density low-stimulation-artifact μLED optoelectrode to the area- and power-efficient CMOS interfacing IC, the presented headstage implements the bidirectional opto-electrophysiology composed with 128 stimulation and 256 recording sites, that is the largest scale reported up to date, with the miniature device size and weight of 23.8 × 28.8 mm^2^ and 3.5-gram, respectively. This eventually results in the highest channel densities in the headstage size (0.56 channels/mm^2^) and weight (109.71 channels/gram) among the state-of-the-art optogenetic systems. The fabricated optogenetic headstage was successfully validated in the *in vivo* experiments observing neuronal interconnections in mouse hippocampus through optical stimulations. All the results offered in this article support that the presented ultrahigh-density, high-channel-count headstage can be a powerful tool to implement the massive-scale optogenetics for comprehensive understanding of the complex deep-brain neuronal networks.

## Methods

### Headstage PCB fabrication

The headstage is implemented on the miniaturized PCB. The headstage PCB consists of 8 layers with 6 dedicated power planes (3 ground layers and 3 power layers) to ensure the robust operation of the CMOS IC by facilitating low resistivity. The conductive area of the PCB was fabricated with 1 oz Cu with 4 mil resolution. The PCB surface was finished by electroless nickel immersion gold (ENIG) for high conductivity and low signal distortion. The gold-plated ground pad was exposed on the PCB surface to bond the CMOS interface IC on the PCB. All the vias on the PCB were tented to avoid a short circuit failure during the device integration. MSOP-8 packaged low dropout regulators (LDOs) (LT3020, Linear Technology, CA, USA), and TSOT-23 packaged voltage references (LT6656, Linear Technology, CA, USA) can be optionally soldered to provide the *in-situ* regulated supply and reference voltages to the CMOS interface IC for less-noisy performance. 1005-size (1.0 mm × 0.5 mm) surface-mounted device (SMD) resistors, capacitors, and ferrite beads are used as passive components for interference decoupling and common mode voltages generation. Transient-voltage-suppression (TVS) diodes with 1005-size package were also integrated to protect the active resistors in the stimulation circuit from electrostatic discharge (ESD) damage.

### Headstage integration

Three miniature data/power connectors (PZN-18-AA, *Omnetics*, MN, USA) were soldered on the bare headstage PCB through the reflow oven. The leads of the connectors were physically fixed by the nonconductive epoxy (353ND-T, *Epoxy Technology*, MA, USA). After cleaning the surface of the PCB using acetone and isopropyl alcohol, the CMOS interface IC was glued on the gold-plated ground pad by applying the nonconductive epoxy (353ND, *Epoxy Technology*, MA, USA) under the chip partially. The fast-drying silver paint (16040-30, *Ted Pella Inc.*, CA, USA) was injected through the ground via holes at the gold-plated ground pad to electrically short the p-substrate of the CMOS interface IC to the PCB ground. The flexible polyimide interposing cable was assembled with the CMOS interface IC through the ball-bump bonding that generates 80 μm diameter gold balls at 100 °C (1^st^-ball-bump bonding). After the 1^st^ bonding, the CMOS interface IC was electrically connected to the headstage PCB through the wirebonding (wedge bonding) that generates 50 μm diameter Al wires at 55 °C. The Al wires and the surface of the CMOS interface IC were covered by the nonconductive epoxy (353ND) for physical protection. The high-density low-stimulation-artifact μLED optoelectrode was attached to the flexible interposing cable through the ball-bump bonding with 80 μm diameter gold balls at 100 °C (2^nd^-ball-bump bonding). The bonding sites on the μLED optoeletrode were covered by the nonconductive epoxy (353ND) for physical protection. The remaining electronic components were additionally soldered on the backside of the headstage PCB.

### Microfabrication of the high-density low-stimulation-artifact μLED optoelectrodes and flexible interposing cable

All the microfabrication steps were carried out at the Lurie Nanofabrication Facility, University of Michigan, Ann Arbor, MI, USA. The procedure for the fabrication of μLED optoelectrodes in a multi-metal-layer configuration [49] was utilized, and the fine-pitch photolithography technique, described in [18], was employed to define narrow metal lines. Throughout the process, a 5× image reduction i-line step-and-repeat projection photolithography tool (GCA AutoStep 200) was utilized for the photolithography step. The fabrication began with a 4” GaN-in-Si LED wafer (Enkris Semiconductor, Suzhou, China) using a low-resistivity boron-doped (111)-Si substrate. After forming mesa LED structures on the LED wafer, interconnecting lines for the LED driving signals were defined, followed by the shielding layer and the electrode signal lines. Metal layers were separated from each other by plasma-enhanced-chemical-vapor-deposited (PECVD) silicon dioxide (SiO_2_) and atomic-layer-deposited (ALD) aluminum oxide (Al_2_O_3_) double-layer passivation. After forming the top passivation, Pt electrode was defined, and the probes were released using a two-step deep reactive ion etching (DRIE) plasma dicing.

The polyimide interposing cable was fabricated in the same microfabrication facility, using a two-mask process similar to the process introduced in [52]. Contact-alignment tool (MA/BA6, Suss Micro Tec SE, Garching, Germany) was used throughout the process. On a 4” silicon wafer with 2 μm thermal SiO_2_ grown on top, 4-μm thick polyimide (PI 2611, HDMicrosystems, Parlin, NJ, USA) layers were formed as the top and the bottom mechanical layers of the cable to be released, and lift-off patterned gold (20 nm chromium, 360 nm gold, 20 nm chromium stack) layer was formed between the top and the bottom polyimide layers to serve as the middle electrical layer. Patterns on the gold layer were designed to have 2-μm half-pitch. After curing the top polyimide layer on the patterned gold layer, oxygen plasma (Plasmatherm 790, Plasma-Therm LLC, Sant Petersburg, FL, USA) was used to etch the whole polyimide outside the cable boundary and inside the via as well as the top polyimide on top of the ball-bumping pads. The cables were mechanically released in buffered hydrofluoric acid (BHF) and thoroughly rinsed in deionized water.

### Design and fabrication of high-channel-count CMOS opto-electrophysiology interface IC

The CMOS interface IC was implemented in the area of 8.7 × 3.7 mm^2^. Except for the pads connected to the μLED optoelectrode (stimulation circuit outputs and recording circuit inputs), all the power and signals were located on the top of the chip for easy wirebonding process. The stimulation circuit outputs and recording circuit inputs were located on the chip core for the direct chip-to-cable integration. The total 424 ball-bump bonding pads with an 80 × 80 μm^2^ size were configured on the chip core with 150 μm pad-to-pad pitch. The test I/O pads were located on the right, left, and bottom sides of the chip for standalone characterizations of the fabricated chip before headstage integration. The CMOS interface IC was fabricated by 180-nm mixed-signal 1P6M CMOS process with 3.3 V I/O supply voltage. For implementing the high-voltage compliance LED driving, the I/O pads associated with the stimulation circuits were customized to exclude the ESD protection cells. The other I/O pads use the standard ESD protection cells provided by the corresponding process design kit.

### Back-end control hardware/software system implementation

The back-end control PCB was implemented with the CPLD microcontroller (Max II, *Intel*, CA, USA). The CPLD serializes the interface IC control commands, and deserializes the acquired neural signals from the recording channels. The high current compliance LDOs (LT3021, Linear Technology, CA, USA) on the back-end control PCB supply the regulated headstage power and the gate voltage of the active resistors used in the stimulator circuits. The data and power are transferred between the headstage and the back-end control PCB through three custom fabricated 18-pin micro cables using polarized miniature connectors (PZN-18-DD, *Omnetics*, MN, USA). The back-end controller PCB is operated with a 9 V battery. The PCI-Express digital I/O device (PCIe-6536, *National Instruments*, TX, USA) is used to interconnect the controller PCB to the custom graphic user interface software programs to adjust the stimulation parameters as well as to monitor recorded neural signals. The software programs were implemented through Python and C.

### *In vivo* experiments and data analysis

All animal handling and experimental procedures associated with or performed in this study followed National Institutes of Health (NIH) animal use guidelines and were approved by the Institutional Animal Care and Use Committee (IACUC) at University of Michigan (Approval Number PRO00009668). An adult male mouse (Thy1-ChR2-YFP, 31 g) was kept in the vivarium on a 12-hour light/dark cycle with ad-libitum access to water and food. On the day of experimentation, the subject was transferred into an anesthetic induction chamber with a constant isoflurane flow at 5 %. The subject was administered a subcutaneous atropine injection (0.05 mg/kg) and secured into a stereotaxic frame with the heater set to 37 C and an isoflurane flow between 1 – 2 % for maintenance. Stages of anesthesia were maintained by confirming the lack of a nociceptive reflex. After confirming the anesthetic plane, the skin head was shaved and cleaned. Immediately after, lidocaine was administered subcutaneously, and a patch of skin was removed to expose the skull. The surface of the skull was cleaned with hydrogen peroxide (2 %). A stainless-steel ground screw was placed above the cerebellum and sealed with dental cement (Unifast LC, GC America). The edges of the craniotomy were marked (-1.25 mm posterior to Bregma and ±1.2 mm lateral to midline) and a 2.4 mm in length craniotomy was drilled. After the dura was removed the device was inserted into the brain to the target depth (-2.2 mm from the surface of the brain).

The device was slowly inserted into the brain to the target depth. The position of the device was confirmed by signals characteristics. The device was set to rest in the target position for 30 minutes before starting the experiment to allow the surrounding tissue to decompress. After recording a 15-minute baseline, LEDs were individually triggered 100 times at 2 Hz and 20 % duty cycle for each power (5, 10 and 20μA). Electrophysiological signals and LED states were sampled at 20 kS/s. Spike sorting was performed using Kilosort [64] and cured in Phy. Cured data was further analyzed and ploted using CellExplorer [65] and custom Matlab scripts. At the end of the experiment the mouse was euthanized with anesthetic overdose and a secondary euthanasia method was implemented as well.

## Author contributions

S.O., K.K., S.P., and E.Y. designed the research. K.K. and E.K. designed and fabricated the high-density low-stimulation-artifact optoelectrode and flexible polyimide interposing cable. S.O., N.S., H.S. and S.P. designed and characterized the high-channel-count CMOS interface IC. S.O. and N.S. designed the headstage PCB and back-end control systems. S.O., N.S., E.K., and V.L. performed the headstage integration. J.R.L.R. and M.V. performed the *in vivo* experiments and analyzed the results. S.O., K.K., J.R.L.R., and S.P. wrote the paper.

---

**Supplementary Fig. 1.**
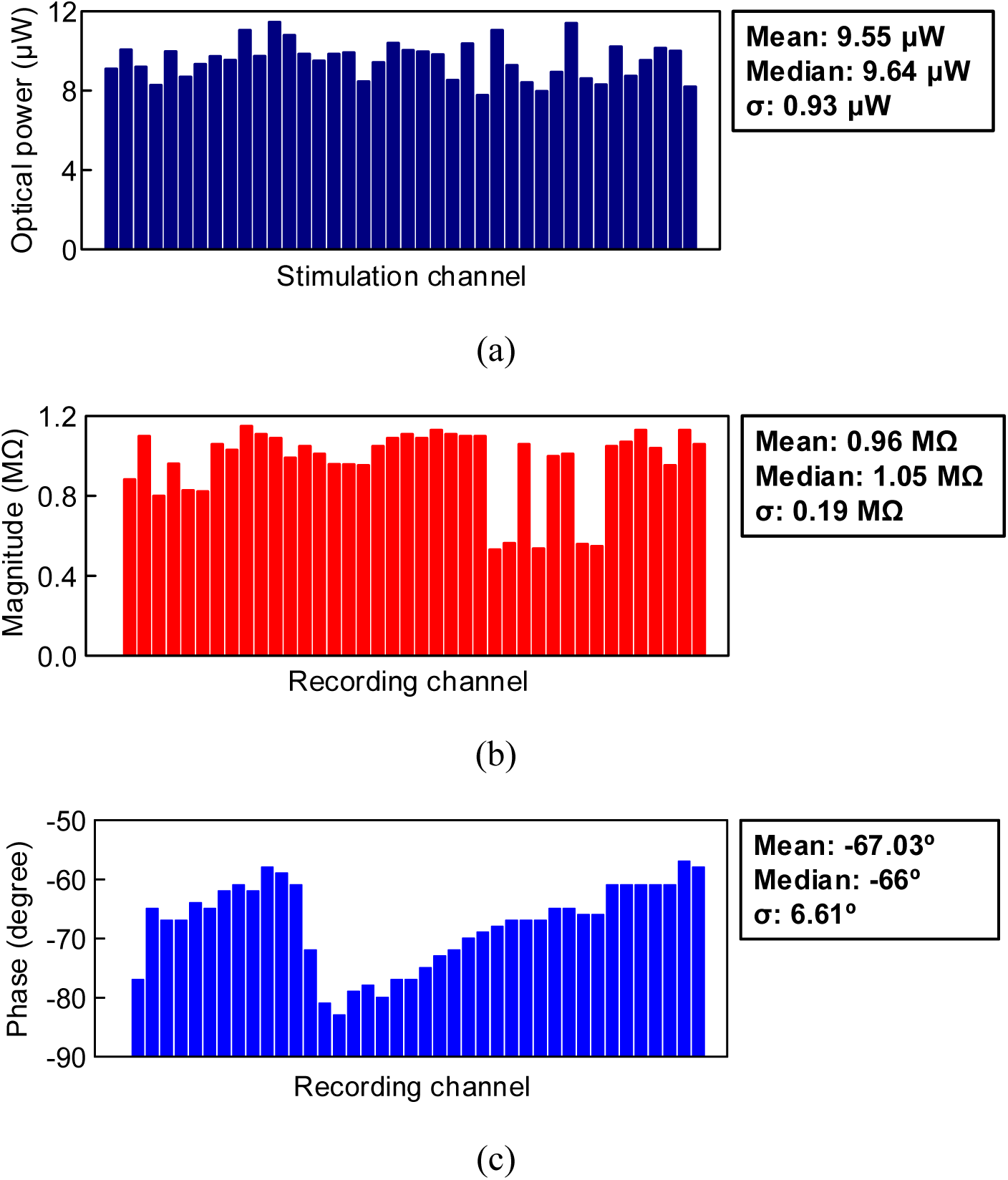
Measured characteristics of the fabricated high-density low-stimulation-artifact μLED optoelectrode. (a) Optical power of the fabricated μLEDs with 100 μA forward current (n = 40). (b) Impedance magnitude of the fabricated Pt recording electrodes at 1 kHz (n = 40). (c) Impedance phase of the fabricated Pt recording electrodes at 1 kHz (n = 40).

**Supplementary Fig. 2.**
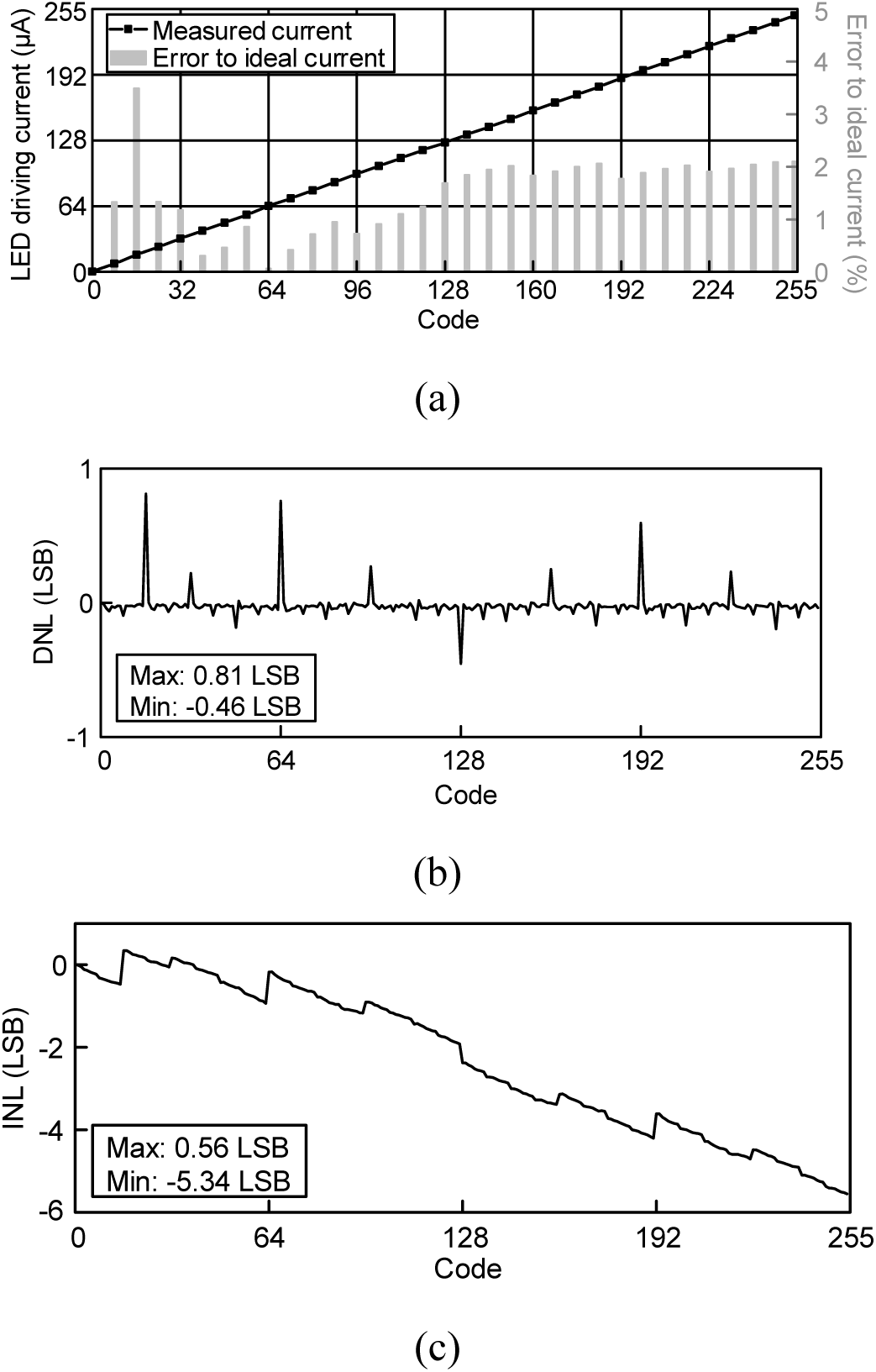
Measured characteristics of the optical stimulation circuit with the coarse-fine output stage design. (a) Measured output current amplitude of the stimulation circuit. (b) Measured differential nonlinearity (DNL) of the current-steering DAC. (c) Measured integral nonlinearity (INL) of the current-steering DAC.

**Supplementary Fig. 3.**
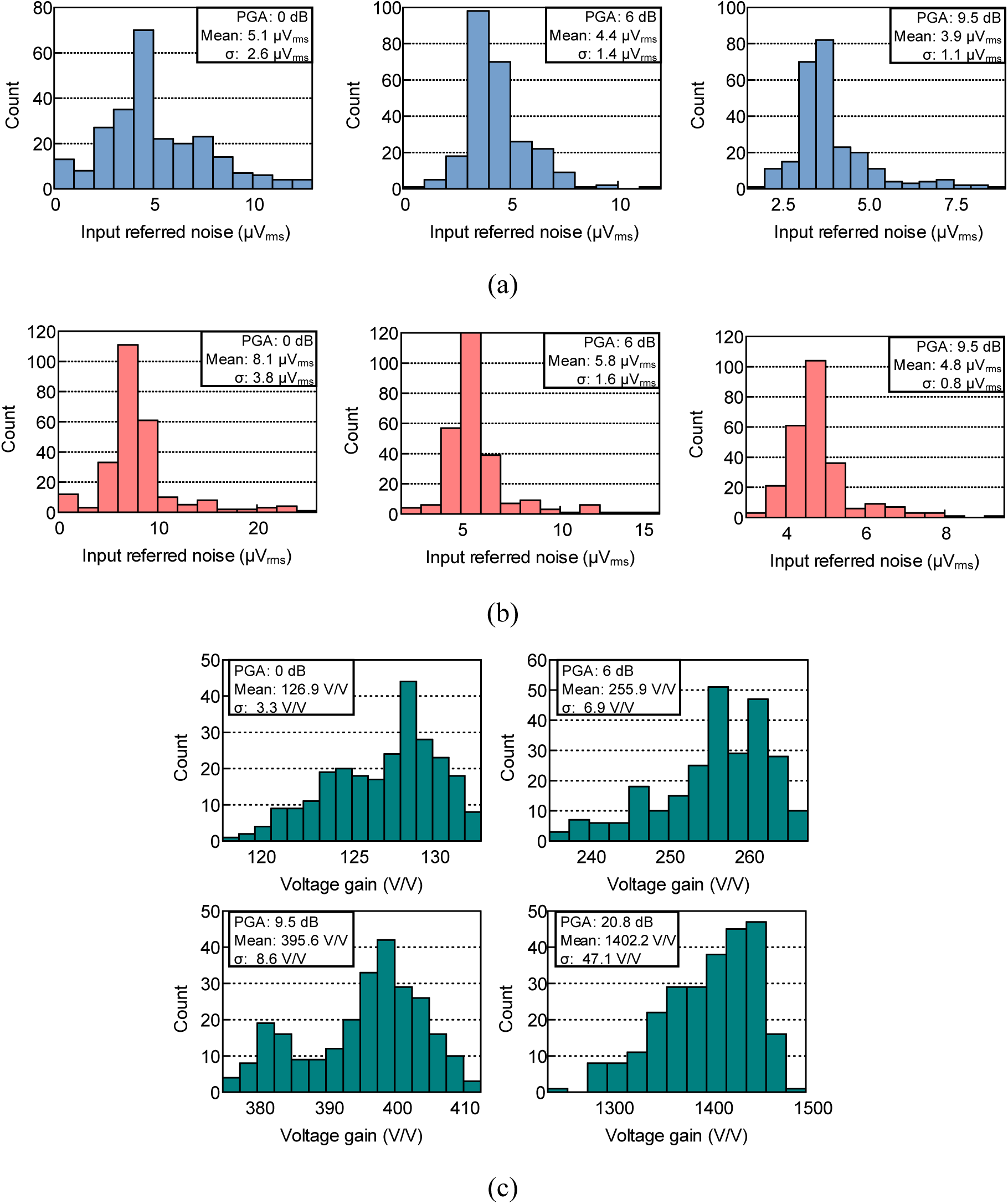
Measured characteristics of the broadband neural recording circuit. (a) Measured input referred noise in LFP band (1 – 300 Hz) with different gain options. (b) Measured input referred noise in AP band (300 Hz – 10 kHz) with different gain options. (c) Measured voltage amplification gains with different gain options.

**Supplementary Fig. 4.**
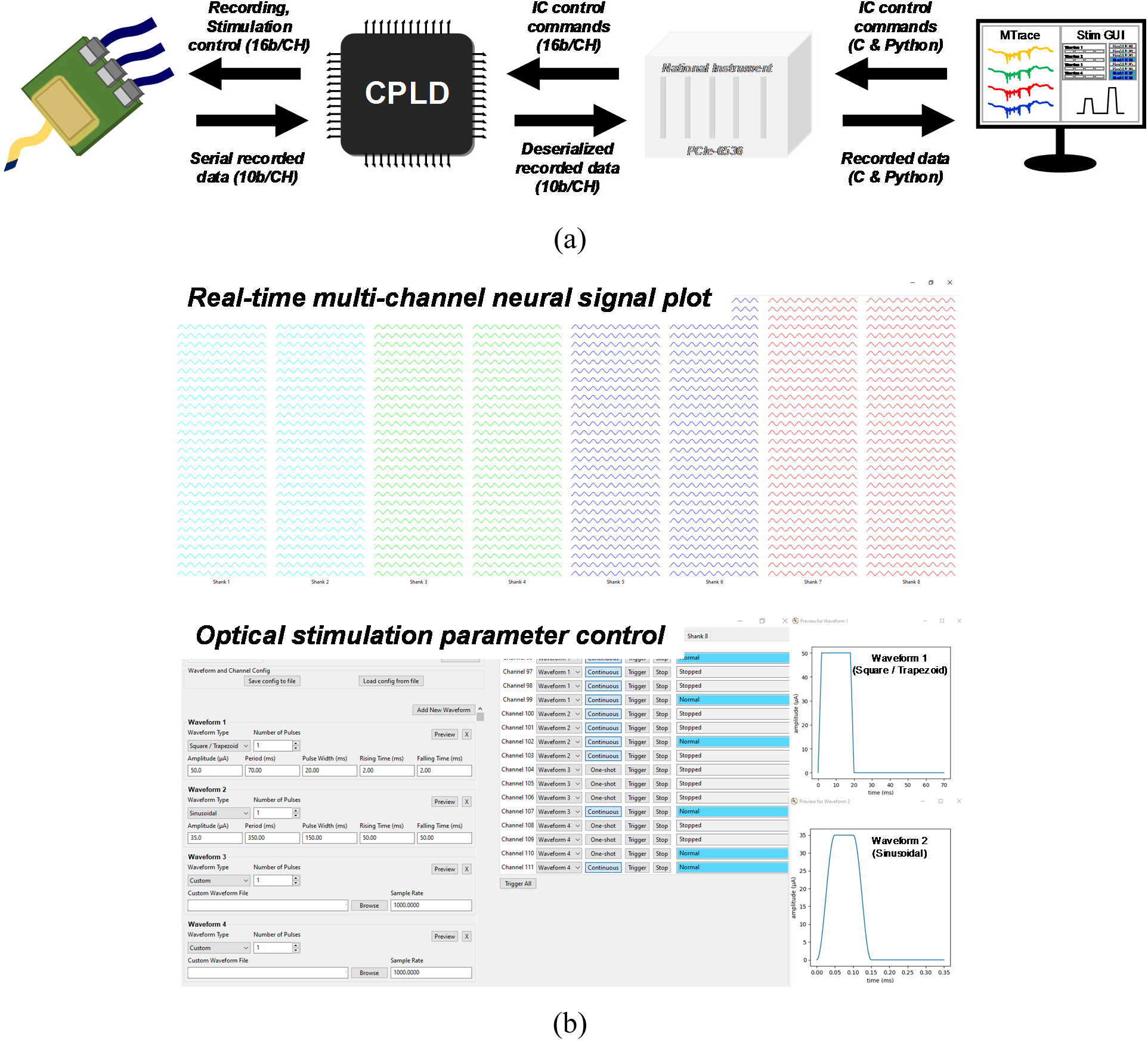
Implementation of the back-end control system. (a) Overall system configuration and signal-chain from the headstage to the software programs. (b) Custom graphic user interface software programs: Real-time waveform plot of the 256-channel neural signals and optical stimulation parameter control of 128 μLEDs.

**Supplementary Fig. 5.**
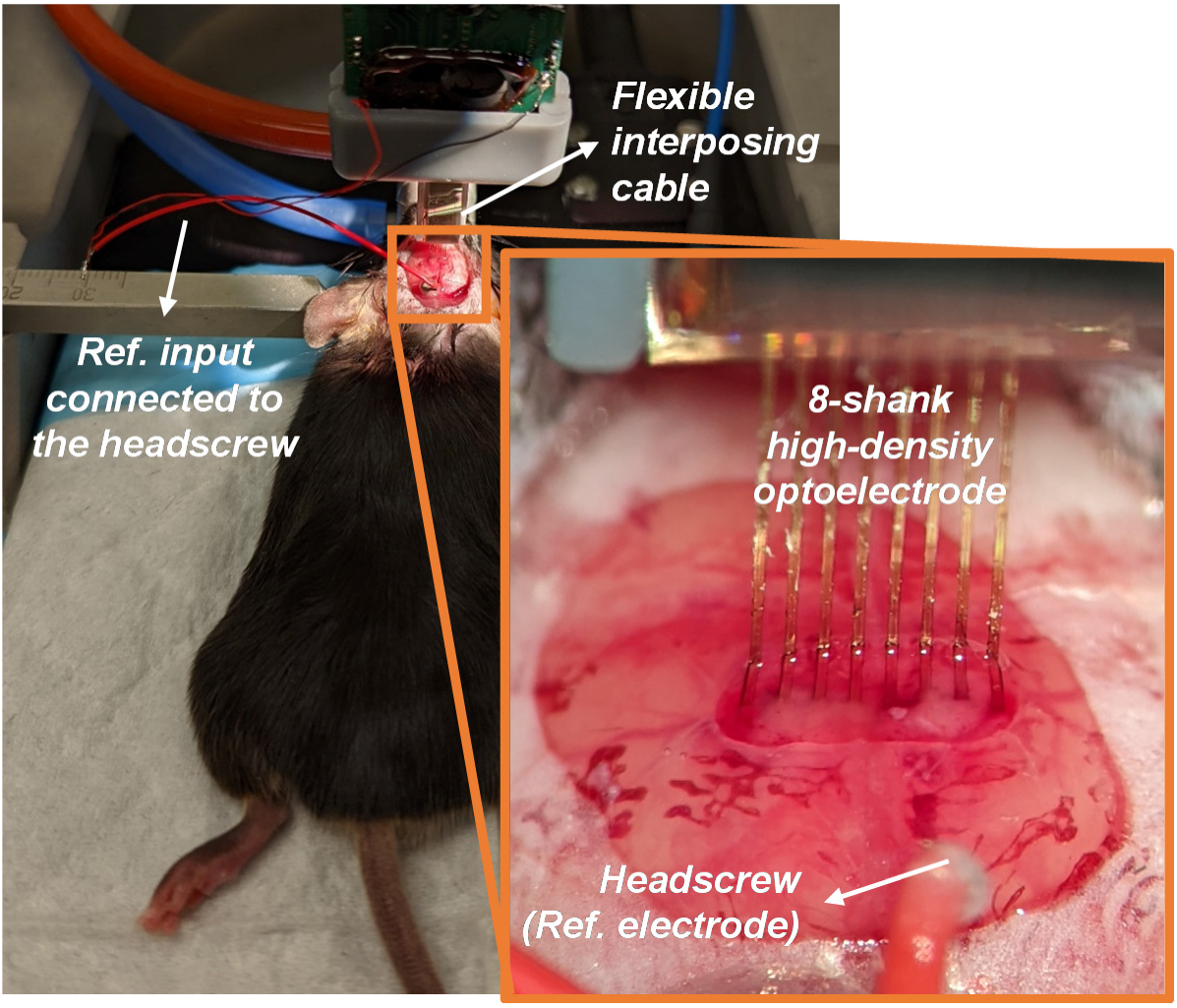
Experimental configuration of the *in vivo* tests.

**Supplementary Fig. 6.**
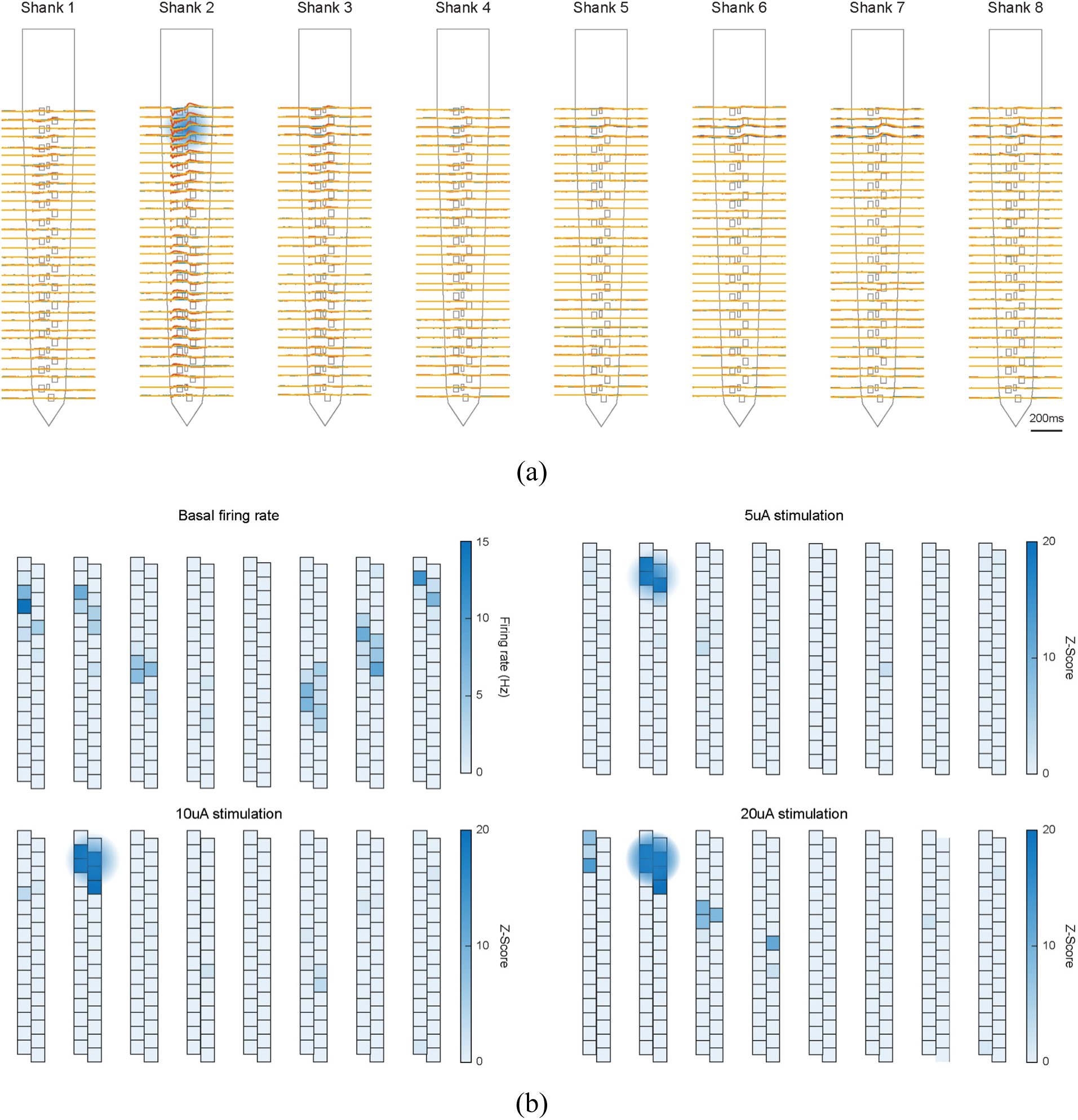
(a) LFP band responses with the stimulation current of 5 µA (sky blue), 10 µA (yellow), and 20 µA (orange): The LFP magnitudes become larger as the stimulation intensity is increased. (b) Firing rate in response to different stimulation intensities: The hippocampal circuitry with pyramidal neurons becomes highly activated as the stimulation intensity is increased.

**Supplementary Fig. 7.**
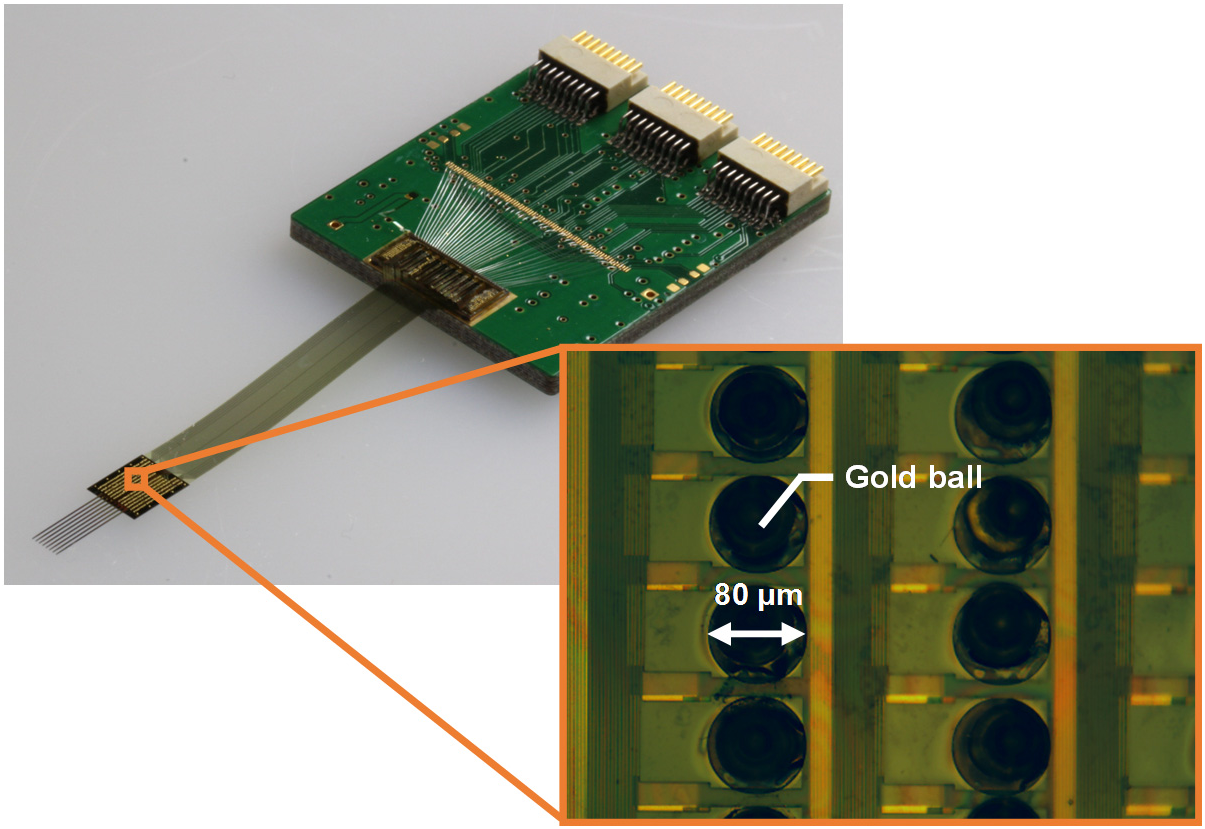
Microphotograph of the ball-bump bonding sites with 80 μm gold balls.

**Supplementary Table 1.** Comparison of the high-density low-stimulation-artifact μLED optoelectrode to the other probes used in the recent optogenetic systems.

|  |  | **This work** | [33] | [34] | [38] | [40] | [59] | [60] | [61] | [62] | [63] |
| --- | --- | --- | --- | --- | --- | --- | --- | --- | --- | --- | --- |
| Probe type |  | **μLED opto electrode** | μLED opto electrode | μLED opto electrode | μLED opto electrode | Optrode w/ opticfiber | μLED needle | μLED needle | μLED needle | Planar LED array | Planar array, Tungsten needle |
| No. of Ch. | Optical stimulation | **128** | 12 | 12 | 12 | 1 | 2 | 1 (4LEDs) | 1, 2, 4 | 16 | 16 |
|  | Other stimulation | - | - | - | - | - | 4 | 1 | - | - | 1 |
|  | Recording | **256** | 32 | 32 | 32 | 8 | - | - | - | - | 16 |
| No. of Shank |  | **8** | 4 | 4 | 4 | 9 | 2 | 1 | 1, 2 | - | 1 |
| Optic source | Type | μLED | μLED | μLED | μLED | Off-the-shelf LED chip | Off-the-shelf LED chip | Off-the-shelf LED chip | Off-the-shelf LED chip | Off-the-shelf LED chip | Off-the-shelf LED chip |
|  | Material | **GaN/InGaN** | GaN/InGaN | GaN | GaN | GaN | InGaN | InGaN | Inorganic | InGaN | InGaN |
|  | Size (μm<sup>2</sup>) | **88** | 160 | 150 | 150 | - | 59400 | 59400 | 59400 | 59400 | 59400 |
|  | Power (mW/mm<sup>2</sup>) | **107.7 @100μA** | 41.1 @100 μA | 33 @100 μA | 80 @0.9mA | 70 @150mA | - | 30 @0.2W | 40 @2.8V | 10 @10mA | 31.5 @24.8mA |
| Recording electrode | Material | **Pt** | Ir | Ir | Ir | Tungsten | - | - | - | - | - |
|  | Size (μm<sup>2</sup>) | **165** | - | - | - | - | - | - | - | - | - |
|  | Impedance (MΩ) | **0.97 @1kHz** | 1 @1kHz | 1.5 @1kHz | 1 @1kHz | 1 | - | - | - | - | - |

**Supplementary Table 2.** Comparison of the high-channel-count CMOS opto-electrophysiology IC to the other interface/controller circuits used in the recent optogenetic systems.

|  |  | **This work** | [33] | [34] | [38] | [40] | [59] | [60] | [61] | [62] | [63] |
| --- | --- | --- | --- | --- | --- | --- | --- | --- | --- | --- | --- |
| Interface/controller circuit |  | **ASIC** | ASIC (Stim)<br>COTS (Rec) | ASIC | ASIC (Stim)<br>COTS (Rec) | COTS | COTS | COTS | COTS | ASIC | ASIC |
| Bidirectional operation |  | **Yes** | Yes | Yes | Yes | Yes | No | No | No | No | Yes |
| No. of Ch. | Optical stimulation | **128** | 48 | 12 | 32 | 32 | 2 | 1 | 1, 2, 4 | 16 | 16 |
|  | Other stimulation | - | - | - | - | - | 4 | 1 | - | - | 8 |
|  | Recording | **256** | 32 | 32 | 32 | 32 | - | - | - | - | 16 |
| Stimulation circuit | Mode | **Current** | Current | Current | Current | Current | Voltage | Voltage | Voltage | Switched-capacitor | Switched-capacitor |
|  | Resolution (bit) | **8** | 10 | 10 | 10 | - | - | - | - | 3 | 5 |
| | Maximum stimulus | **255 μA** | 1.023 mA | 1 mA | 900 μA | 100 mA | 2.7–3.0 V | - | 2.8 V | 10 mA | 24.8 mA |
| Recording circuit | Bandwidth (kHz) | **10** | 10 | 15 | 10 | 10 | - | - | - | - | 10 |
|  | Resolution (bit) | **10** | 16 | 10 | 16 | 16 | - | - | - | - | 10 |
| | Noise (μV<sub>rms</sub>) | **5.4** | 2.4 | 5.7 | 2.4 | 2.4 | - | - | - | - | 3.46 |
| Chip size (mm<sup>2</sup>) |  | **30.71** | 68.5 | 8.13 | 65.4 | 96 | 88.16 | 27 | 1.45 (1ch),<br>17.81 (2ch),<br>19.49 (4ch) | 1.1 | 15 |
| Size/channel (mm<sup>2</sup>) |  | **0.08** | 0.86 | 0.18 | 1.02 | 1.5 | 14.69 | 13.5 | 1.45 (1ch),<br>8.91 (2ch),<br>4.87 (4ch) | 0.07 | 0.38 |

